# Thyroid hormone receptor beta inhibits the PI3K-Akt-mTOR signaling axis in anaplastic thyroid cancer via genomic mechanisms

**DOI:** 10.1101/2020.11.12.379933

**Authors:** Cole D. Davidson, Eric L. Bolf, Noelle E. Gillis, Lauren M. Cozzens, Jennifer A. Tomczak, Frances E. Carr

**Author notes:** **Cole D. Davidson**: Conceptualization, Formal analysis, Investigation, Writing – Original draft preparation, Visualization. **Eric L. Bolf:**Conceptualization, Formal analysis, Investigation, Writing – Review & editing. **Noelle E. Gillis**: Formal analysis, Investigation, Writing – Review & editing. **Lauren M. Cozzens:**Formal analysis, Investigation. **Jennifer A. Tomczak:**Investigation, Writing – Review & editing. **Frances E. Carr:**Resources, Writing – Review & editing, Supervision, Project administration, Funding acquisition.

## Abstract

Thyroid cancer is the most common endocrine malignancy, and the global incidence has increased rapidly over the past few decades. Anaplastic thyroid cancer (ATC) is highly aggressive, dedifferentiated, and patients have a median survival of fewer than six months. Oncogenic alterations in ATC include aberrant PI3K signaling through receptor tyrosine kinase (RTK) amplification, loss of phosphoinositide phosphatase expression and function, and Akt amplification. Furthermore, the loss of expression of the tumor suppressor thyroid hormone receptor beta (TRβ) is strongly associated with ATC. TRβ is known to suppress PI3K in follicular thyroid cancer and breast cancer by binding to the PI3K regulatory subunit p85α. However, the role of TRβ in suppressing PI3K signaling in ATC is not completely delineated. Here we report that TRβ indeed suppresses PI3K signaling in ATC through unreported genomic mechanisms including a decrease in RTK expression and increase in phosphoinositide and Akt phosphatase expression. Furthermore, the reintroduction and activation of TRβ in ATC enables an increase in the efficacy of the competitive PI3K inhibitors LY294002 and buparlisib on cell viability, migration, and suppression of PI3K signaling. These findings not only uncover additional tumor suppressor mechanisms of TRβ but shed light into the implication of TRβ status and activation on inhibitor efficacy in ATC tumors.

Graphical abstract

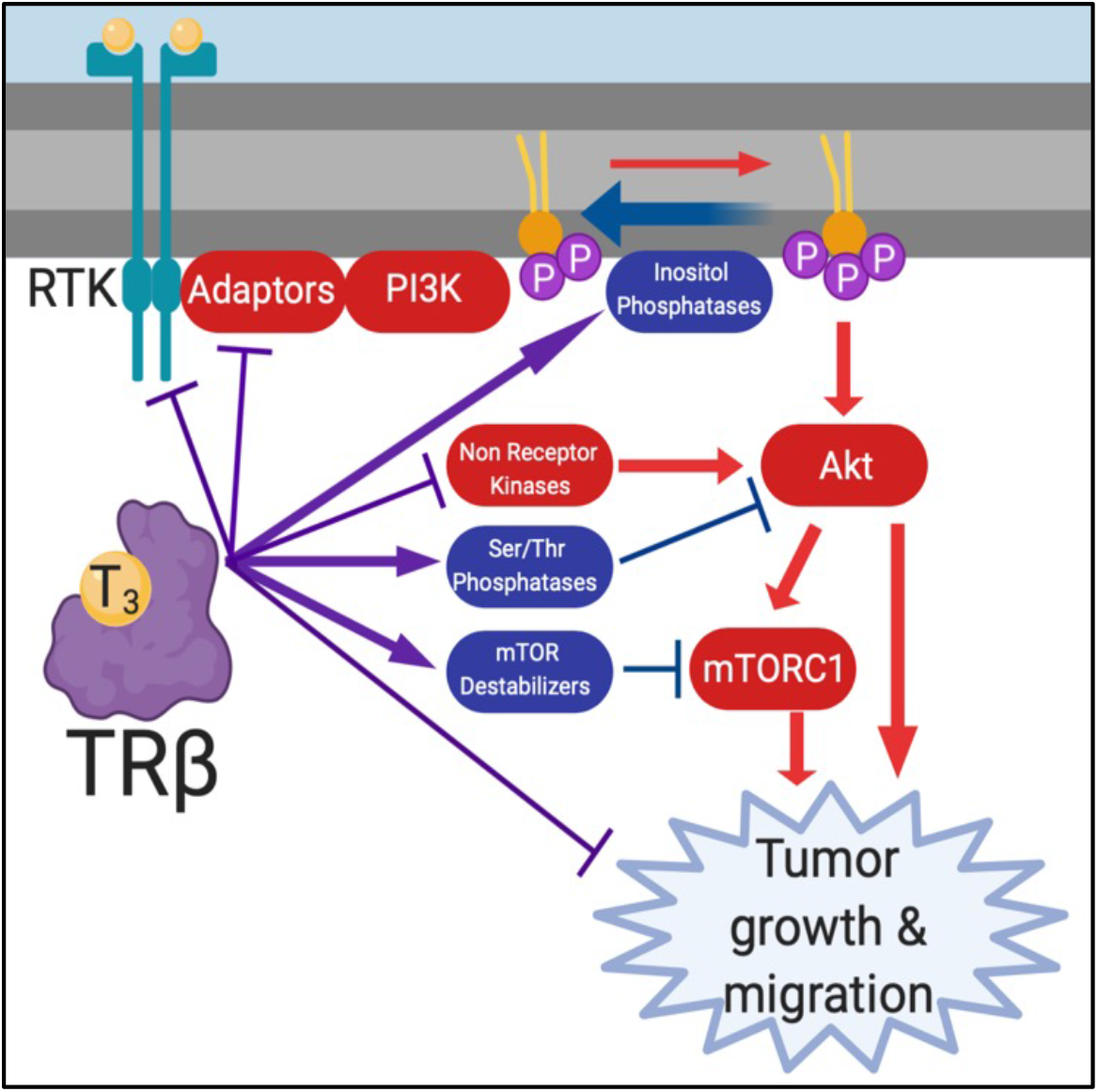

## 1. Introduction

Thyroid cancer is the most common endocrine malignancy, and the incidence has been rapidly increasing the past few decades (1). While the overall prognosis for thyroid cancer is generally favorable, patients with the most aggressive and dedifferentiated subtype, anaplastic thyroid cancer (ATC), have a median survival of three to five months (2). The current most effective treatments for ATC patients increase survival time by only a few months as drug resistance and tumor recurrence often develop (3). Therefore, there is an unmet need for more precise understanding of the molecular etiology of ATC tumorigenesis and new strategies for improving patient outcome.

Phosphoinositide 3 kinase (PI3K) signaling is a prominent molecular driver for aggressive and poorly differentiated thyroid cancers such as ATC (4). PI3K is recruited to the plasma membrane by phosphorylated, ligand-bound receptor tyrosine kinases (RTKs). PI3K phosphorylates the membrane lipid phosphatidylinositol 4,5-bisphosphate (PI(4,5)P_2_) to PIP_3_, which recruits 3-phosphoinositide dependent protein kinase 1 (PDPK1) and the mammalian target of rapamycin complex 2 (mTORC2) to the plasma membrane to phosphorylate Akt on thr308 and ser473, respectively. Phosphatase and tensin homolog (PTEN) is a tumor suppressor that dephosphorylates PIP_3_ back to PI(4,5)P_2_. Akt phosphorylates a myriad of targets that are involved in cell cycle progression and survival signaling. In addition, Akt leads to the activation of mTORC1, which phosphorylates targets such as p70S6K that ultimately lead to the activation and assembly of translation factors for protein synthesis and cell growth (5,6). Multiple genes are either mutated or amplified within the PI3K pathway in ATC. Frequent alterations include amplification of RTKs such as epidermal growth factor (EGFR), amplification or gain-of-function mutations in *PIK3CA* (PI3K), loss-of-function mutations or decreased expression of *PTEN*, and amplification of *AKT1* (7–10).

In addition to the canonical mechanisms of PI3K regulation, there are a multitude of other factors that regulate the pathway. These factors include members of the nuclear hormone receptor family including estrogen, androgen, and thyroid hormone receptors (11–13). Our work has demonstrated that thyroid hormone receptor beta (TRβ) acts as a tumor suppressor in ATC through repression of several pathways important for tumor growth (14,15). However, the potential for TRβ to exhibit tumor suppression in ATC by suppressing PI3K is not fully understood. Multiple groups have reported the potential for TRβ to bind to the regulatory subunit of PI3K, p85α, to inhibit phosphorylation of PI(4,5)P_2_ to PIP_3_ (16–18). While these mechanisms help explain the potential for TRβ to inhibit PI3K in certain cancer models, there may be other mechanisms of TRβ-mediated suppression. While TRβ has been shown to inhibit PI3K via nongenomic mechanisms in breast (19) and follicular thyroid (20,21) cancer, it is unknown if this mechanism or unexplored genomic mechanisms occur in ATC. Therefore, we sought to better understand the mechanism of TRβ-mediated suppression of PI3K signaling in ATC using a TRβ-expression model. Moreover, we tested the efficacy of the PI3K inhibitors LY294002 and buparlisib in cells with or without TRβ expression. These findings present previously unexplored mechanisms of the tumor suppression by TRβ, the role of TRβ in ATC, as well as the implication of TRβ expression status in response to PI3K-targeted therapeutic intervention.

## 2. Materials and methods

### 2.1. Culture of thyroid cell lines

Cells were cultured in RPMI 1640 growth media with L-glutamine (300 mg/L), sodium pyruvate and nonessential amino acids (1%) (Corning Inc, Corning, NY, USA), supplemented with 10% fetal bovine serum (Thermo Fisher Scientific, Waltham, MA, USA) and penicillin-streptomycin (200 IU/L) (Corning) at 37°C, 5% CO2, and 100% humidity. Lentivirally modified SW1736 cells were generated as described (15,22) with either an empty vector (SW-EV) or to overexpress TRβ (SW-TRβ). SW-EV and SW-TRβ were grown in the above conditions with the addition of 2 μg/ml puromycin (Gold Bio, St Louis, MO, USA). SW1736 and KTC-2 were authenticated by the Vermont Integrative Genomics Resource at the University of Vermont (Burlington, Vermont) using short tandem repeat profiles and Promega GenePrint10 System (SW1736, May 2019; KTC-2, October 2019). 8505C, OCUT-2, and CUTC60 were authenticated by University of Colorado by short tandem repeat profiles (8505C, June 2013; OCUT-2, June 2018; CUTC60, November 2018).

### 2.2. Cell culture reagents

Triiodothyronine (T_3_) was purchased from Sigma (St. Louis, MO, USA) and dissolved in 1 N NaOH and diluted to 10 nM in cell culture medium at the time of each application. LY294002 and buparlisib were purchased from MedChemExpress (Monmouth Junction, NJ, USA). LY294002 was dissolved in 100% ethanol and buparlisib was dissolved in 100% DMSO prior to indicated dilutions for cell culture experiments.

### 2.3. Immunoblot analysis

Proteins were isolated from whole cells in lysis buffer (20mM Tris-HCl (pH 8), 137mM NaCl, 10% glycerol, 1% Triton X-100, and 2mM EDTA) containing Protease Inhibitor Cocktail (Catalogue Number 78410) (Thermo Fisher Scientific), 1 mM Na_3_VO_4_, and 1 mM PMSF (Sigma). Proteins were quantified via Pierce Coomassie Plus (Bradford) Assay (Thermo Fisher Scientific), and 25 μg of protein per sample were resolved by polyacrylamide gel electrophoresis on 10% Tris-Glycine gels (Catalogue Number XP00105BOX) (Thermo Fisher Scientific) and immobilized onto nitrocellulose membranes (GE Healthcare, Chicago, IL, USA) by electroblot (Bio-Rad Laboratories, Hercules, CA, USA). Membranes were blocked with 5% w/v BSA in TBS and 0.1% v/v Tween20 (Gold Bio, St Louis, MO, USA) for one hour at room temperature and incubated with primary antibodies overnight (Supplementary Table 1); immunoreactive proteins were detected by enhanced chemiluminescence (Thermo Scientific) on a ChemiDoc XRS+ (Bio-Rad Laboratories). Densitometry analysis in Figure 4B was performed using ImageJ (NIH, Bethesda, MD, USA).

### 2.4. Measurement of Akt phosphorylation by ELISA

Akt serine 473 phosphorylation was measured using Pathscan phospho-Akt1 sandwich ELISA kit, according to the manufacturer's instructions (Cell Signaling Technology, Danvers, MA, USA). Samples were prepared from cells treated with 10 nM T_3_ for 24 hours and then 1 hour incubation in the presence or absence of 1 or 10 μM LY294002. 100 μl of samples containing equal amount of protein were applied to each well.

### 2.5. Measurement of PI3K activity by ELISA

PI3K activity was determined using a commercially available PI3K ELISA kit (Echelon Biosciences Inc., Salt Lake City, UT) according to the manufacturer's instructions. Briefly, after drug treatment, cells were washed in ice-cold PBS and lysed in 500 μl ice-cold lysis buffer (137 mM NaCl, 20 mM Tris–HCl (pH 7.4), 1 mM CaCl_2_, 1 mM MgCl_2_, 1 mM Na_3_VO_4_, 1% NP-40 and 1 mM PMSF). PI3K was then immunoprecipitated with 5 μl of antibody (anti-p85α, MilliporeSigma, Burlington, MA, USA) and 60 μl of Pierce Protein A/G magnetic beads (Thermo Scientific). PI3K activity in the immunoprecipitates was then assayed by PI3K ELISA according to the manufacturer's instructions. The spectrophotometric data were obtained using a Synergy 2 Multi-Detection Microplate Reader (Agilent Technologies, Santa Clara, CA, USA) at a wavelength of 450 nm. The protein concentrations of cellular lysates were determined by Bradford assay as described above. The activity of PI3K was corrected for protein content.

### 2.6. RNA-seq data analysis of PI3K pathway intermediates

Previously published RNA-seq data was used to determine expression levels of genes within the PI3K pathway (15). Construction of the PI3K signaling genes of interest for our study was based on the curated IPA pathway gene set. Additional genes of interest were added based on literature search and cancer relevance. Normalized transcript counts generated with DESeq2 were used to calculate fold change compared to the control condition (SW-EV-T_3_). Raw and processed expression data can be found in the Gene Expression Omnibus (GEO) database under accession number GSE150364.

### 2.7. Analysis of thyroid cancer patient sample data

Publicly available microarray expression data, deposited in the GEO Database (GSE76039, GSE3467, GSE82208), were analyzed using GEOR2 (www.ncbi.nlm.nih.gov/gds) to reveal differential expression of genes relevant to PI3K signaling across the spectrum of thyroid cancers.

### 2.8. RNA extraction and quantitative Real-Time PCR (qRT-PCR)

Total RNA was extracted using RNeasy Plus Kit (Qiagen) according to manufacturer’s protocol. cDNA was then generated using the 5X RT Mastermix (ABM, Vancouver, Canada). Gene expression to validate RNA-seq analysis was quantified by qRT-PCR using BrightGreen 2X qPCR MasterMix (ABM, Vancouver, Canada) on a QuantStudio 3 real-time PCR system (Thermo Fisher Scientific). Fold change in gene expression compared to endogenous controls was calculated using the ddCT method. Primer sequences are indicated in Supplemental Table 2.

### 2.9. In vitro cell viability assay

The cell viability assay was performed by plating 1.0 × 10^4^ SW-EV or SW-TRβ cells into 12 well (22.1 mm) tissue culture dishes. After adhering overnight, the cells were treated with 10 nM T_3_ and LY294002, buparlisib, or vehicle at the indicated concentrations. Every day after treatment for four days, the medium was removed, cells were washed with PBS and lifted with trypsin (Thermo Scientific), and the number of surviving cells was counted with a hemocytometer.

### 2.10. Migration assay

Cell migration was determined by wound healing assay. Cells were plated and allowed to grow to 100% confluency. Two hours prior to scratching, cells were treated with 10 μg/ml Mitomycin C (Sigma, MO, USA) dissolved in H_2_O. A scratch was performed with a P1000 pipette tip and debris was washed away with PBS. Migration media was supplemented with 10 nM T_3_ and LY294002 or vehicle. Images were obtained using a Canon digital camera connected to an Axiovert inverted microscope (Carl Zeiss, Oberkochen, Germany) at 0, 24, 48, and 72 hours. Wound closure was measured using ImageJ macro “Wound Healing Tool” (http://dev.mri.cnrs.fr/projects/imagej-macros/wiki/Wound_Healing_Tool). Values were normalized so that the initial scratch was 0% closure.

### 2.11. Statistics

All statistical analyses were performed using GraphPad Prism software. Paired comparisons were conducted by Student’s t-test. Group comparisons were made by one-way ANOVA followed by Dunnett’s or Tukey’s multiple comparison test as appropriate. Two-way ANOVA followed by a Tukey’s multiple comparison test w conducted for multigroup analysis. Data are represented as mean ± standard deviation. Area under the curve (AUC) at the 95th confidence interval was used to evaluate statistical differences in growth and migration assays.

## 3. Results

### 3.1 Rapid thyroid hormone receptor action fails to suppress PI3K in ATC cells

TRβ is a known suppressor of the PI3K signaling pathway in breast and follicular thyroid cancer. This has been previously described as a nongenomic mechanism in which TRβ binds to the regulatory subunit of PI3K, p85α, preventing recruitment to ligand-bound RTKs (13). This action is rapid, and the addition of the thyroid hormone triiodothyronine (T_3_) modulates the response within 15-30 minutes (23). Therefore, we sought to evaluate the impact of short term T_3_ treatment in the SW1736 cell line with restored stable expression of TRβ or an empty vector (EV) control. Analysis of pAkt and pmTOR by Western blot surprisingly revealed a minimal impact of TRβ with or without T_3_ on PI3K suppression (Fig. 1A). To validate these results, we performed a PI3K immunoprecipitation followed by ELISA to test the ability of PI3K to catalyze the phosphorylation of PI(4,5)P_2_ to PIP_3_ following T_3_ treatment. Again, we observed a modest but insignificant decrease in PIP_3_ production in the presence of TRβ and T_3_ (Fig. 1B).

**Figure 1.**
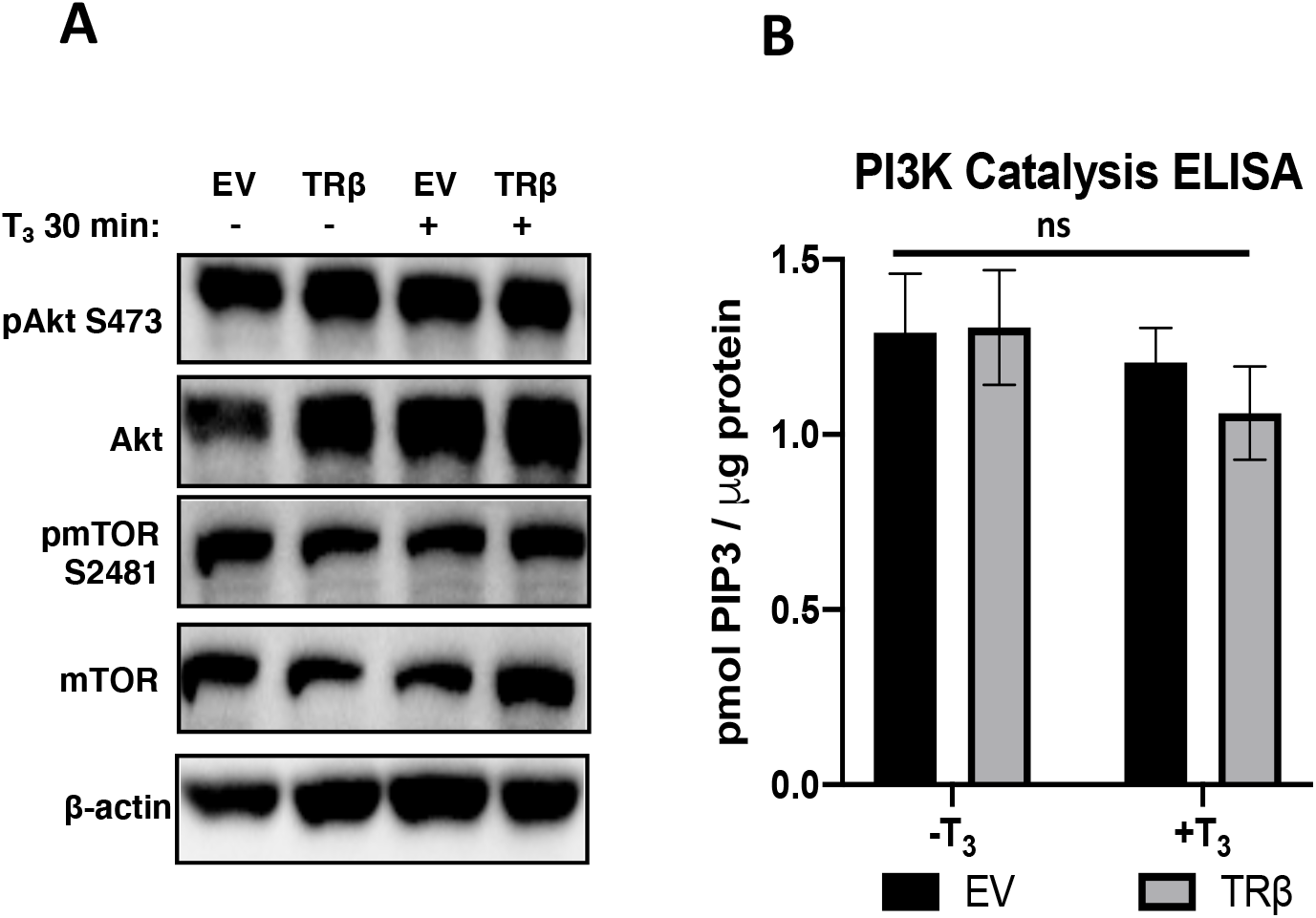
Short-term or no exposure to T_3_ is insufficient for TRβ-mediated PI3K suppression. SW1736-EV (EV) and SW1736-TRβ (TRβ) cells were treated with 10 nM T_3_ or vehicle (1 N NaOH) for 30 min before protein levels were determined by immunoblot (**A**) or incubated with anti-p85α antibody for PI3K catalysis ELISA (**B**). ELISA signal in **B** was standardized to protein concentration as determined by a Bradford assay. Significance in **B** was calculated by two-way ANOVA followed by Tukey multiple comparisons test. ns = no significance (p ≥ 0.05) across treatment groups.

### 3.2 Long-term T_3_ treatment suppresses PI3K signaling in ATC cells

Since short-term T_3_ treatment did not suppress PI3K activity in our cell line model, we hypothesized that long-term T_3_ treatment may enable TRβ-mediated suppression of PI3K signaling. Therefore, we treated our EV and TRβ cells with T_3_ for 24 hours then measured pAkt, pmTOR, and pp70S6K levels by Western blot (Fig. 2A). The SW-TRβ cells treated with T_3_ exhibited a marked decrease in pAkt on serine 473 but not threonine 308. Serine 473 phosphorylation induces a substantial increase in Akt activity following growth factor stimulation and plays a role in regulating substrate specificity (24). Additionally, there was a modest reduction in pP70S6K, a kinase further downstream of Akt. To validate the observed Akt ser473 dephosphorylation, we conducted a sandwich ELISA with our EV and TRβ cells following 24-hour T_3_ treatment (Fig. 2B); pAkt (ser473) was again reduced with liganded-TRβ.

**Figure 2:**
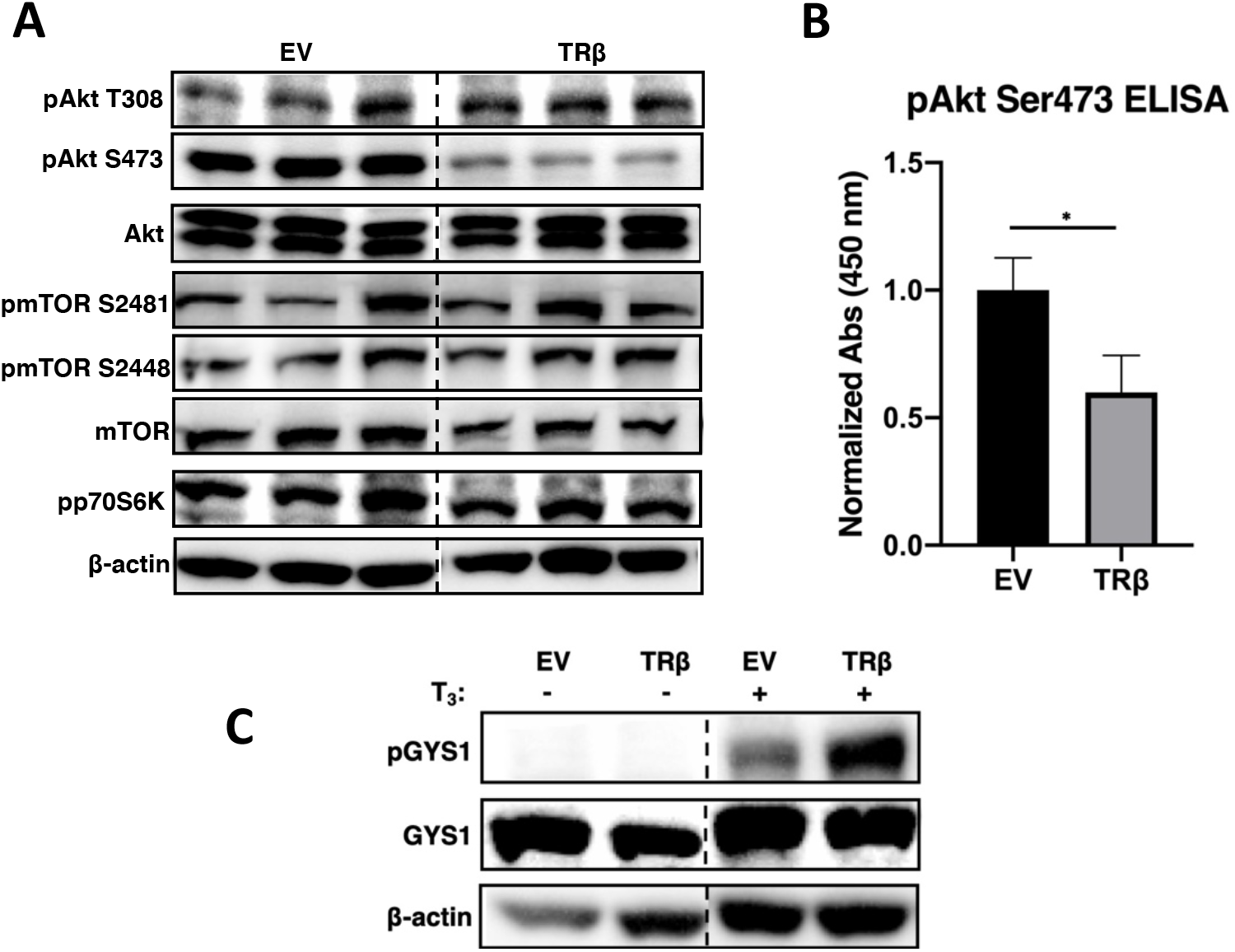
Long-term T_3_ with TRβ inhibits the PI3K-Akt-mTOR pathway in SW1736 cells. SW1736-EV (EV) and SW1736-TRβ (TRβ) cells were treated with 10 nM T_3_ for 24 hours before protein levels were determined by immunoblot (**A** and **C**) or subjugated to a pAkt Ser473 sandwich ELISA (**B**). The samples in **A** are replicates. Dashed lines in immunoblots indicate gap between two sets of lanes on the same membrane. ELISA signal in **B** was standardized to protein concentration as determined by a Bradford assay. Significance in **B** was calculated by Student’s t-test. (*p < 0.05).

In order to test the impact of long-term T_3_ exposure and heightened TRβ expression we measured a well-studied downstream effector, glycogen synthase kinase 3 beta (GSK3β) and its substrate glycogen synthase 1 (GYS1). GSK3β is a multi-substrate kinase and, importantly, is implicated in the progression of numerous cancers including ATC (25,26). We observed a modest increase in phosphorylated, and thus inactivated, GYS1 in the presence of T_3_ and a marked increase of pGYS1 in the presence of both TRβ and T_3_ (Fig. 2C). Since TRβ-T_3_ treatment suppressed both Akt-mTOR-P70S6K and Akt-GSK3β-GYS1 pathways, these data are suggestive of a requirement for both thyroid hormone and TRβ to achieve robust inactivation of PI3K signaling as observed through two separate pathways downstream of Akt.

### 3.3 Liganded-TRβ transcriptionally remodels the PI3K signaling landscape

Since a long-term T_3_ treatment was needed for robust suppression of PI3K signaling in our cell line model, we hypothesized that TRβ may be regulating the expression of key components of this pathway. In order to better understand the extent to which TRβ suppresses PI3K signaling through genomic mechanisms, we leveraged our RNA-sequencing data performed on our EV and TRβ cell lines following 24 hours of T_3_ treatment (15). Numerous genes involved in the PI3K pathway were determined to be differentially expressed in the presence of both T_3_ and TRβ (Fig. 3A-C).

**Figure 3:**
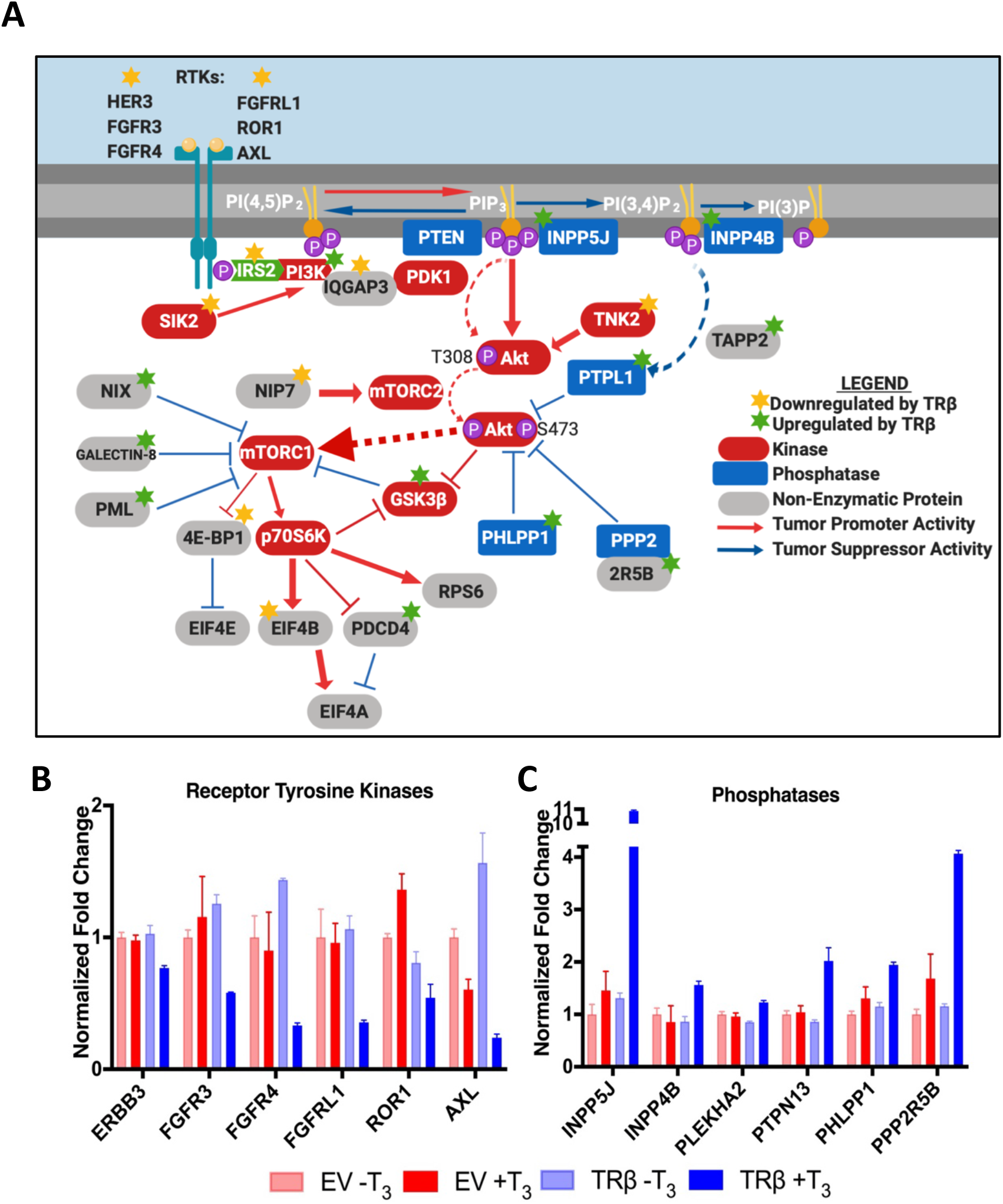
Liganded-TRβ decreases expression of oncogenic genes and increases tumor suppressive genes in the PI3K-Akt-mTOR signaling axis. **A.** RTKs dimerize in response to ligand to allow for IRS2 to dock and recruit PI3K (45). PI3K phosphorylates PI(4,5)P_2_ to PIP_3_. PTEN and INPP5J dephosphorylate PIP_3_ to PI(4,5)P_2_ and PI(3,4)P_2_ respectively. INPP4B dephosphorylates PI(3,4)P_2_ to PI(3)P. PIP_3_ recruits PDK1 (not shown) and Akt to the plasma membrane for PDK1 phosphorylation of Akt T308. NIP7-activated mTORC2 phosphorylates Akt S473, which is dephosphorylated by PHLPP1 (46). ING5 has been shown to dephosphorylate Akt in hormone-dependent cancers (47). The 2R5B subunit of PPP2 directs the complex to dephosphorylate Akt T308 and S473 (30). TNK2 phosphorylates Akt Y176 to enhance plasma membrane recruitment (48). Akt leads to the activation of mTORC1 by TSC 1/2-Rheb (not shown). Nix and PML destabilize Rheb-mTORC1 binding (49). Galectin-8 inhibits and delocalizes mTORC1 (50). GSK3β is inhibited by Akt and inhibits multiple substrates relevant to glucose and glycogen metabolism, survival signaling, and cell cycle progression. mTORC1 phosphorylates and inhibits 4E-BP1 which inhibits EIF4E. MTORC1 also activates P70S6K which inhibits GSK3β and PDCD4 and activates EIF4B and RPS6. EIF4B and PDCD4 regulate EIF4A (49). **B.** Ligand-bound TRβ decreased expression of receptor tyrosine kinases. **C.** Ligand-bound TRβ increased expression of PI3K-Akt phosphatases. Significance and fold change values between EV+T_3_ and TRβ+T_3_ are located in Supplementary Table 1.

Receptor tyrosine kinase gene expression was reduced in the TRβ-T_3_ group, including HER3 (*ERBB3*), fibroblast growth factor receptor isoforms (*FGFR3, FGFR4, FGFRL1*), neurotrophic tyrosine kinase receptor-related 1 (*ROR1*), and tyrosine-protein kinase receptor UFO (*AXL*). While *PTEN* expression was not increased, other membrane-bound phosphoinositide phosphatases did increase including phosphatidylinositol 4,5-bisphosphate 5-phosphatase A (*INPP5J*), and inositol polyphosphate 4-phosphatase type II (*INPP4B*), which dephosphorylate PIP_3_ to PI(3,4)P_2_ and PI(3,4)P_2_ to PI(3)P, respectively (27). There was also an increase in expression of cytosolic phosphatases that dephosphorylate Akt primarily on ser473, including tandem-PH-domain-containing protein-2 and protein-tyrosine-phosphatase-like protein-1 (*PLEKHA2* and *PTPN13*), the R5B localization subunit of protein phosphatase 2 (*PPP2R5B*), and PH domain and leucine-rich repeat-protein phosphatase 1 (*PHLPP1*) (28–30). In addition to these genes that serve to regulate PI3K activation and subsequent Akt phosphorylation, genes involved in PI3K recruitment and stabilization, mTORC regulation, and translation factors were found to be differentially regulated in the TRβ-T_3_ group (Supplemental Fig. 1A-D and Table 3). The fold change of a subset of the differentially expressed genes were validated via RT-qPCR (Supplemental Fig. 2).

### 3.4 Endogenous TRβ expression in ATC cells correlates with low pAkt ser 473 and high PI3K-Akt signaling phosphatase expression

Following our RNA-sequencing findings in transduced SW1736 cells, we next questioned if level of endogenous TRβ expression correlates with reduced Akt phosphorylation and increased PI3K-Akt phosphatase expression. We demonstrated a significant inverse correlation between TRβ expression and Akt phosphorylation as shown by immunoblot (Fig. 4A and B). Furthermore, 8505C with the highest level of endogenous TRβ also had the highest expression of the phosphoinositide phosphatases *INPP4B* and *INPP5J* as well as the pAkt ser473 phosphatase *PHLLP1* (Fig. 4C). These findings illustrate the trend between TRβ and expression of tumor suppressive genes in PI3K signaling in a genetically diverse set of ATC cell lines (Supplemental Table 4).

**Figure 4:**
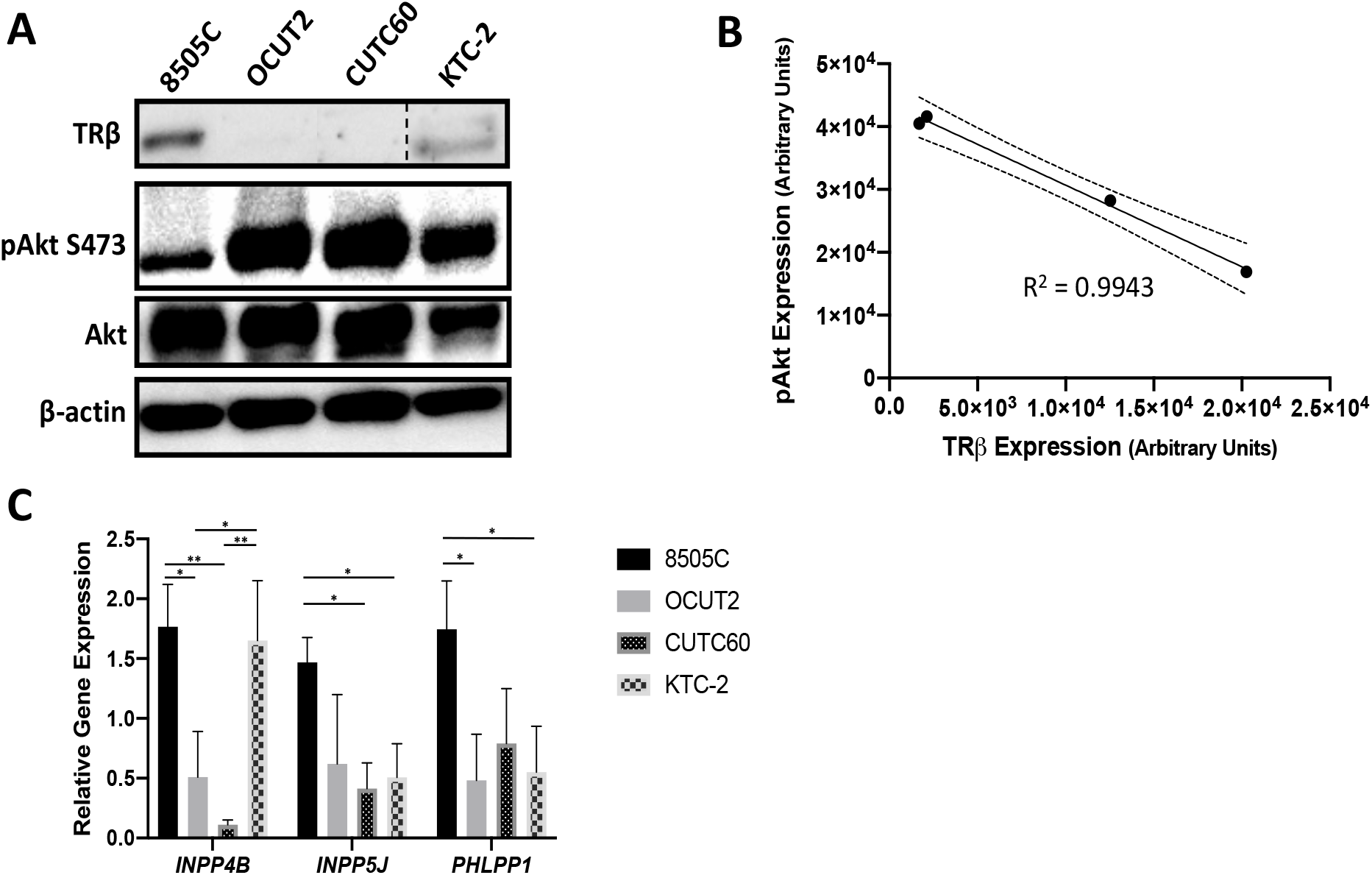
TRβ expression correlates with low Akt ser473 phosphorylation and high phosphoinositide and Akt phosphatase expression. ATC cell lines were analyzed for TRβ and pAkt ser473 expression (**A and B**) and phosphoinositide and Akt phosphatases (**C**). Dashed line in immunoblot indicates gap between two sets of lanes on the same membrane. Significance was determined by one-way ANOVA followed by Dunnett’s multiple comparisons test (* p < 0.05, ** p < 0.01).

### 3.5 PI3K signaling genes regulated by TRβ in SW1736 cells are aberrantly expressed in patient thyroid cancer samples

Next, we used expression data from matched normal tissue and papillary, follicular, poorly differentiated, and anaplastic thyroid carcinomas to determine if any of the PI3K regulators we revealed to be altered by TRβ-T_3_ treatment exhibited differential expression in different thyroid cancer subtypes (31–33). The patient microarray data revealed that the RTKs ERBB3 (HER3), ROR1, and AXL expression levels correlate with TC subtype where expression is highest in the more aggressive tumors (Fig. 5A). Conversely, phosphatase expression is highest in matched normal tissue and differentiated thyroid tumors (Fig. 5B). Interestingly, expression of FGFR4 and FGFRL1 were lowest in the more aggressive PDTC and ATC populations. We next analyzed gene expression data for TRβ and markers of differentiated thyroid cells to demonstrate the connection between TRβ-T_3_ presence with increased RTK and decreased phosphatase expression. *THRB* (TRβ) and genes encoding enzymes and transporters for thyroid hormone synthesis were coordinately lost in FTC, PDTC, and ATC patient samples (Fig. 5C). As demonstrated previously, TRβ protein is significantly reduced in FTC, PDTC, and ATC patients (14), a finding that may contribute to the gene expression data presented here.

**Figure 5:**
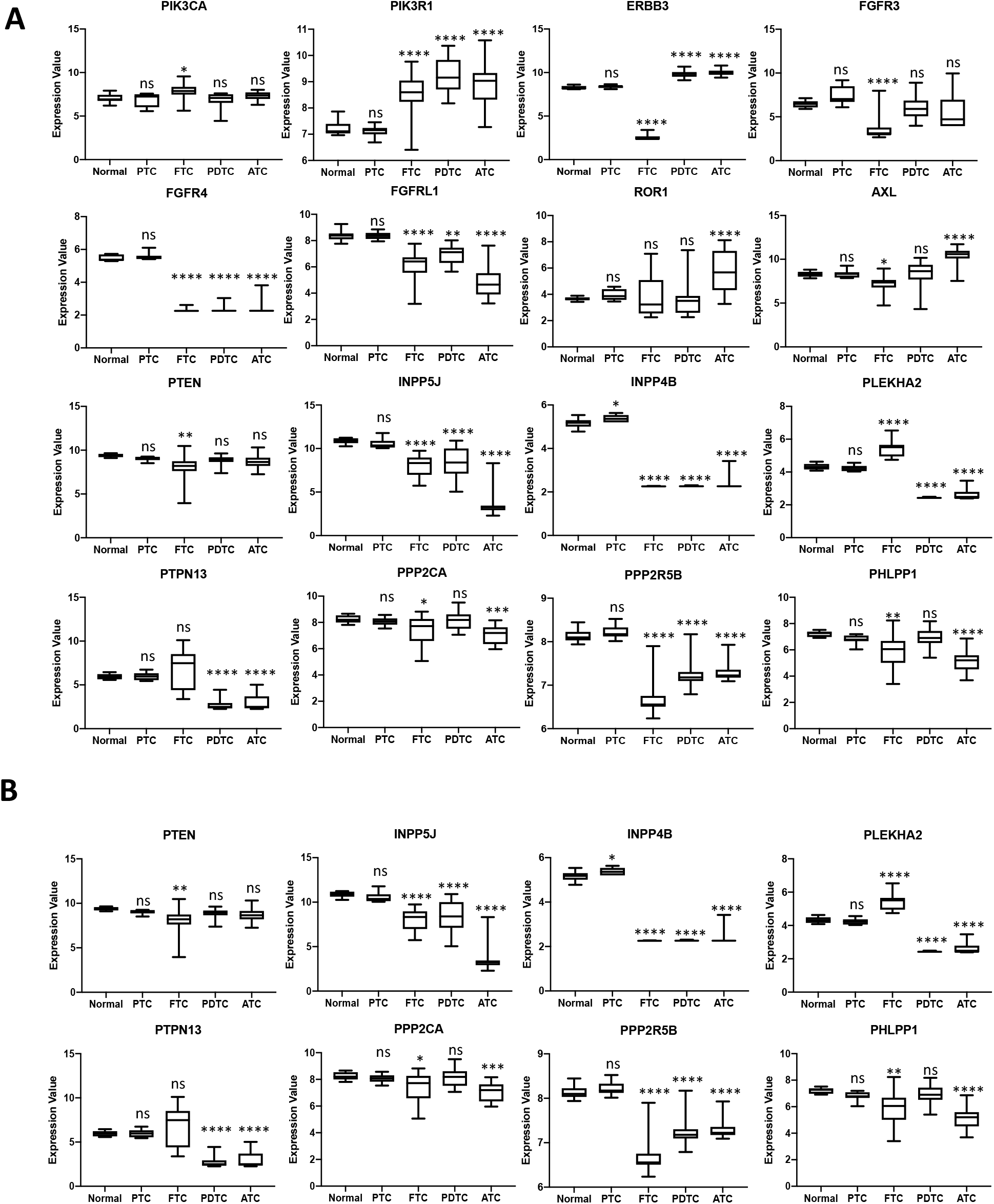
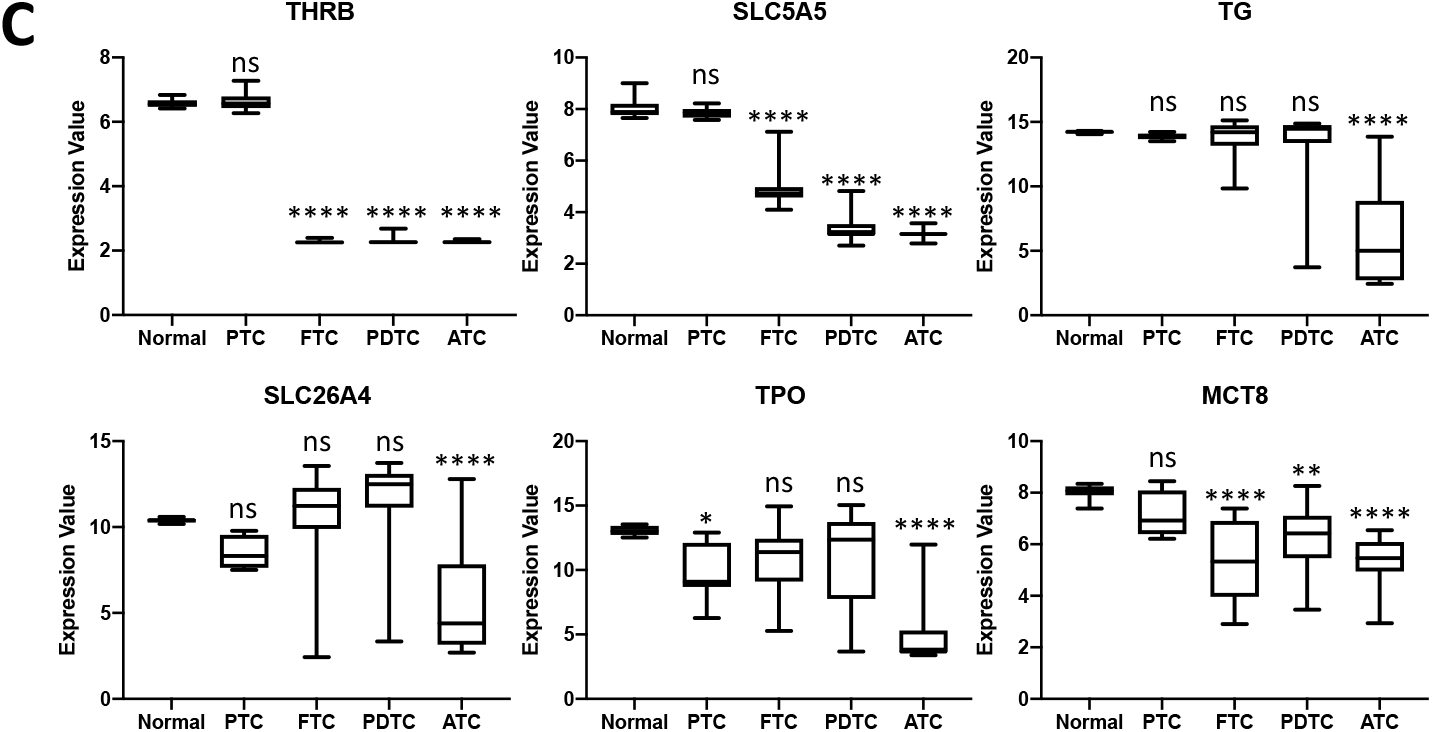
PI3K signaling genes regulated by TRβ in SW1736 cells are aberrantly expressed in patient thyroid cancer samples. Patient thyroid cancer microarray data (GSE76039, GSE3467, GSE82208) were analyzed for genes encoding receptor tyrosine kinases (**A**), phosphoinositide and Akt phosphatases (**B**), and TRβ, enzymes, and transporters necessary for synthesizing thyroid hormones. Significance was determined by one-way ANOVA followed by Dunnett’s multiple comparisons test (ns = no significance p ≥ 0.05, * p < 0.05, ** p < 0.01, ***p < 0.001, **** p < 0.0001). Normal n = 9. PTC n = 9, FTC n = 27, PDTC n = 17, ATC n = 20

### 3.6 TRβ and T_3_ inhibit cell viability and migration in serum-activated cells

PI3K signaling fuels cancer progression by stimulating cell survival, proliferation, and migration. We previously demonstrated that TRβ inhibits SW1736 proliferation in charcoal-stripped media, an observation that was dependent upon T_3_ stimulation (15). To achieve phenotypic confirmation of our sequencing results, we challenged our engineered cells in full serum (10%) media with or without additional 10 nM T_3_ to measure a functional consequence of PI3K inhibition. Even in the presence of full serum and activated RTKs, the TRβ group rendered the SW1736 cells less viable (Fig. 6A and B). In accordance with the RNA-sequencing data, T_3_ is necessary for maximum inhibition of cell viability by TRβ. Additionally, we challenged our cells to migrate in the presence of liganded-TRβ. TRβ cells were unable to effectively migrate to close the wound compared to the EV cells (Fig. 6C-E). These findings are similar to what we observed previously in experiments using charcoal-stripped, growth-deprived media, thus indicating the absolute requirement of TRβ to inhibit cell viability and migration.

**Figure 6:**
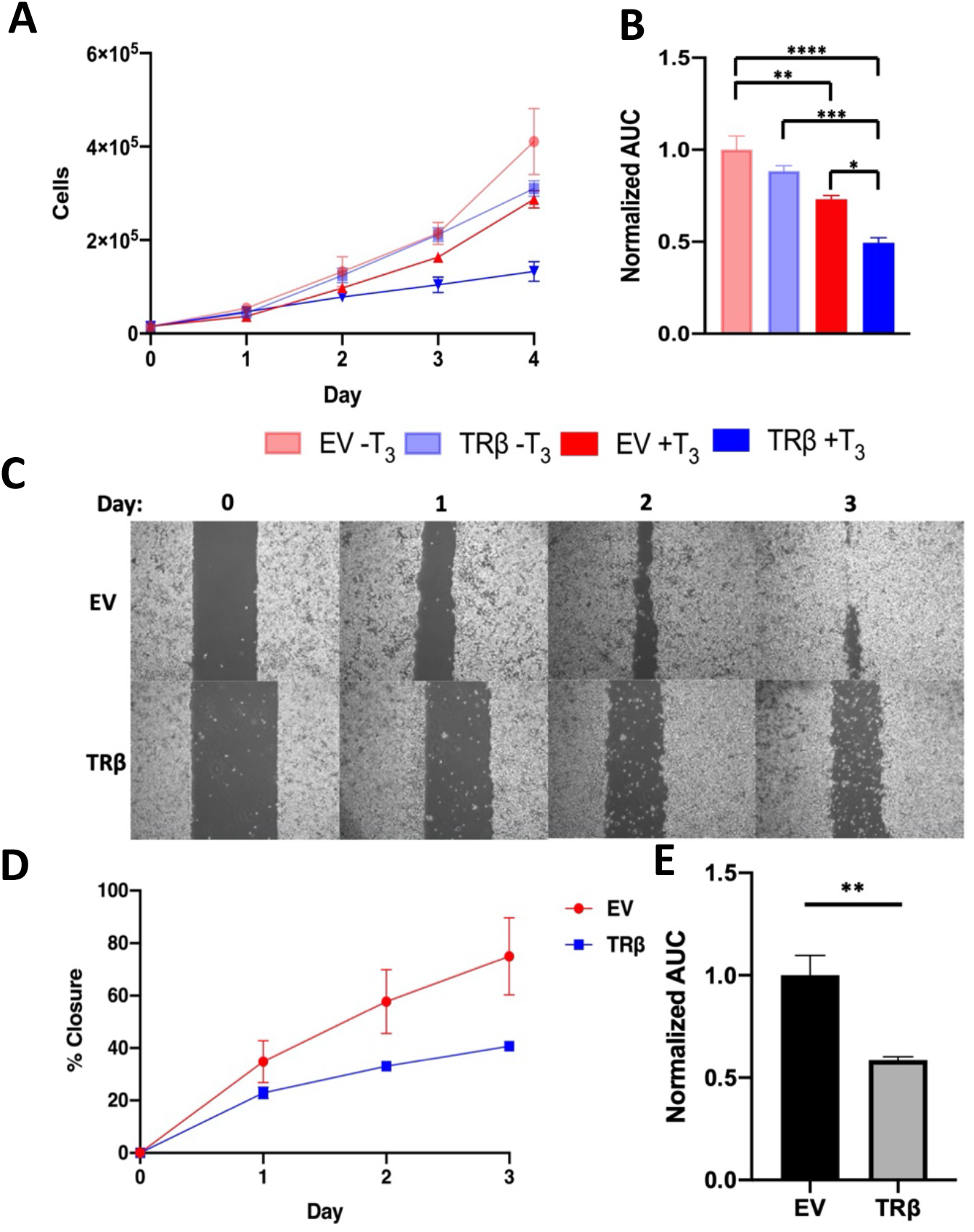
TRβ and T_3_ reduce cell viability and migration in serum-activated SW1736 cells. **A.** SW1736-EV (EV) and SW1736-TRβ (TRβ) cells were treated with 10 nM T_3_ or vehicle (1 N NaOH) for four days. Each day the cells were lifted with trypsin and counted using a hemocytometer for viable cells. Area under the curve (AUC) analysis was performed (**B**) and normalized to the EV-T_3_ group. Significance was calculated using one-way ANOVA followed by Tukey’s multiple comparisons test (* p < 0.05, ** p < 0.01, ***p < 0.001, **** p < 0.0001). **C.** EV and TRβ cells were grown to confluency in 6 well plates before treatment with 10 μg/ml mitomycin C for 2.5 hours. Media were aspirated, and the cells were scratched with a P1000 pipette tip before being washed with PBS and treated with media containing 10 nM T_3_. **D.** Wells were imaged each day and % wound closure relative to day 1 was calculated. **E.** Area under the curve (AUC) analysis was performed and normalized to the EV group. Significance was calculated using Student’s t-test test. ** p < 0.01.

### 3.7 TRβ improves PI3K inhibitor efficacy

In order to further test our hypothesis that TRβ suppresses PI3K signaling in ATC cells, we established the efficacy of small molecule competitive PI3K inhibitors on cell cytotoxicity. Both LY294002 and buparlisib compete for the ATP-binding site of PI3K, preventing PIP_2_ phosphorylation. It should be noted that buparlisib is 27 times more potent than LY294002 (IC50 of 52 nM and 1.4 μM, respectively (34)). The TRβ cells showed an improved response to LY294002, with an EC50 value nearly 5-fold less than the EV cells (Fig. 7A). In addition, the TRβ cells were nearly 25-fold more sensitive to the PI3K inhibitor buparlisib than the EV cells (Supplemental Fig. 3). In concordance with the RNA sequencing and cell viability data, TRβ requires T_3_ to fully exert the tumor suppressive profile; LY294002 efficacy was not increased in SW-TRβ cells lacking 10 nM T_3_ (Supplemental Fig. 4). LY294002 efficacy was also enhanced in the TRβ cells as measured by migration assay at both 1 and 10 μM (Fig. 7B-D).

**Figure 7:**
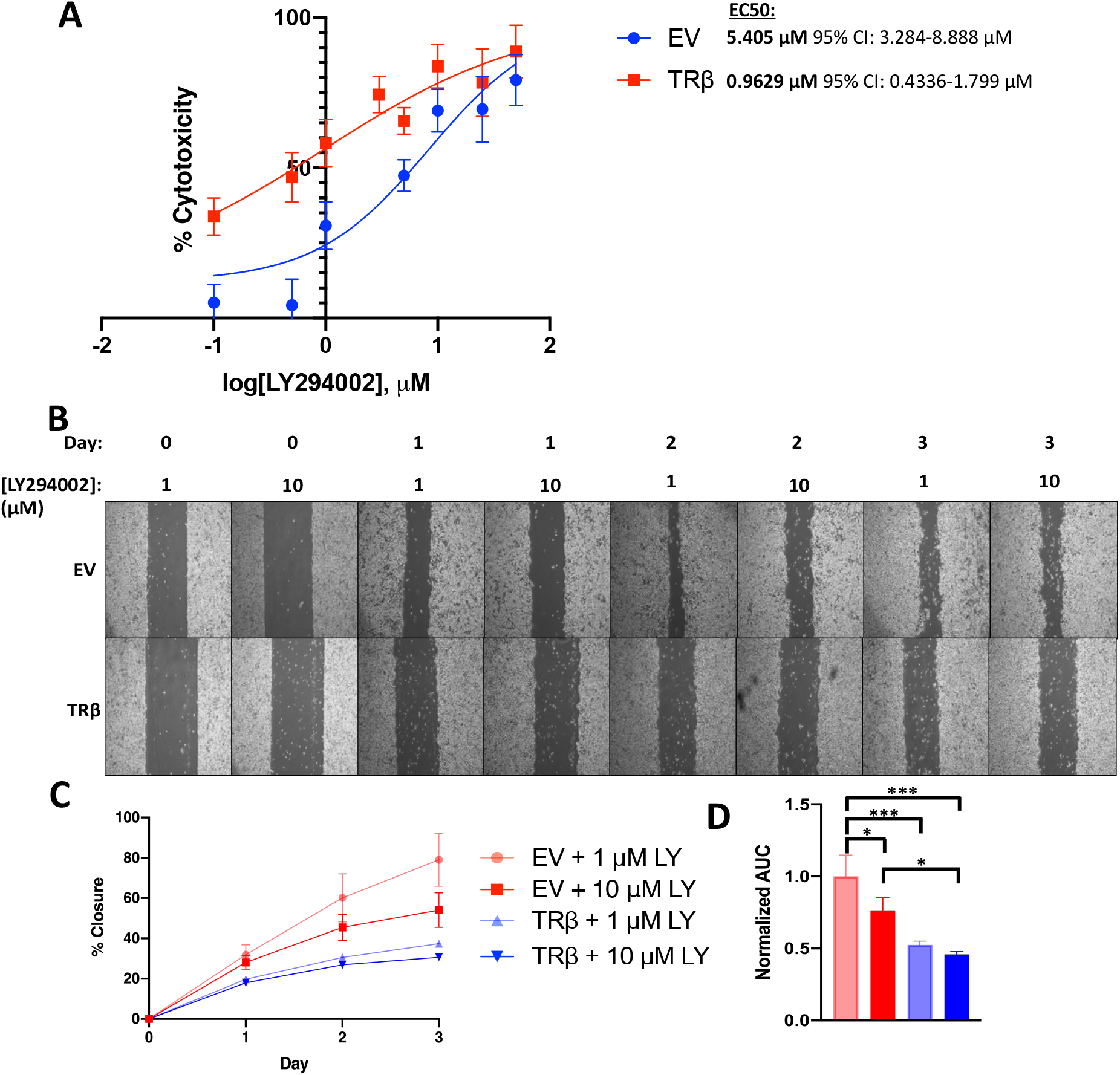
The effect of LY294002 on cytotoxicity and migration is enhanced in SW1736-TRβ cells. **A.** EV and TRβ cells were treated with 10 nM T_3_ and LY294002 (0.1-50 μM) or matched-concentration vehicle (100% EtOH) for four days. Each day the cells were lifted with trypsin and counted using a hemocytometer for viable cells. Area the curve analysis was conducted for each LY294002 concentration to calculate % cytotoxicity relative to EV vehicle at each concentration of LY294002. EC50 values were calculated using GraphPad Prism’s nonlinear regression package. **B.** EV and TRβ cells were grown to confluency in 6 well plates before treatment with 10 μg/ml mitomycin C for 2.5 hours. Media were aspirated, and the cells were scratched with a P1000 pipette tip before being washed with PBS and treated with media containing 10 nM T_3_ and 1 or 10 μM LY294002. **C.** Wells were imaged each day and % wound closure relative to day 1 was calculated. **D.** Area under the curve (AUC) analysis was performed and normalized to the EV 1 μM LY294002 group. Significance was calculated using one-way ANOVA followed by Tukey’s multiple comparisons test (* p < 0.05, ** p < 0.01, ***p < 0.001).

### 3.8 TRβ enhances PI3K inhibitor inactivation of the PI3K-Akt-mTOR axis

To further confirm that TRβ is specifically inhibiting the PI3K signaling pathway, we challenged our cells with LY294002 and buparlisib and measured pAkt, pmTOR, and pp70S6K levels following 24 hours of T_3_ treatment. LY294002 and buparlisib were able to blunt Akt thr308 phosphorylation in both cell lines, but there was a marked decrease in levels of pAkt (ser473), pmTOR (ser2448 and ser2481), and pp70S6K (Fig. 8A and Supplemental Fig. 5A). We validated these findings with a pAkt (ser473) sandwich ELISA and detected lower levels of pAkt in the LY294002-treated TRβ cells compared to the EV control (Figure 8B).

**Figure 8:**
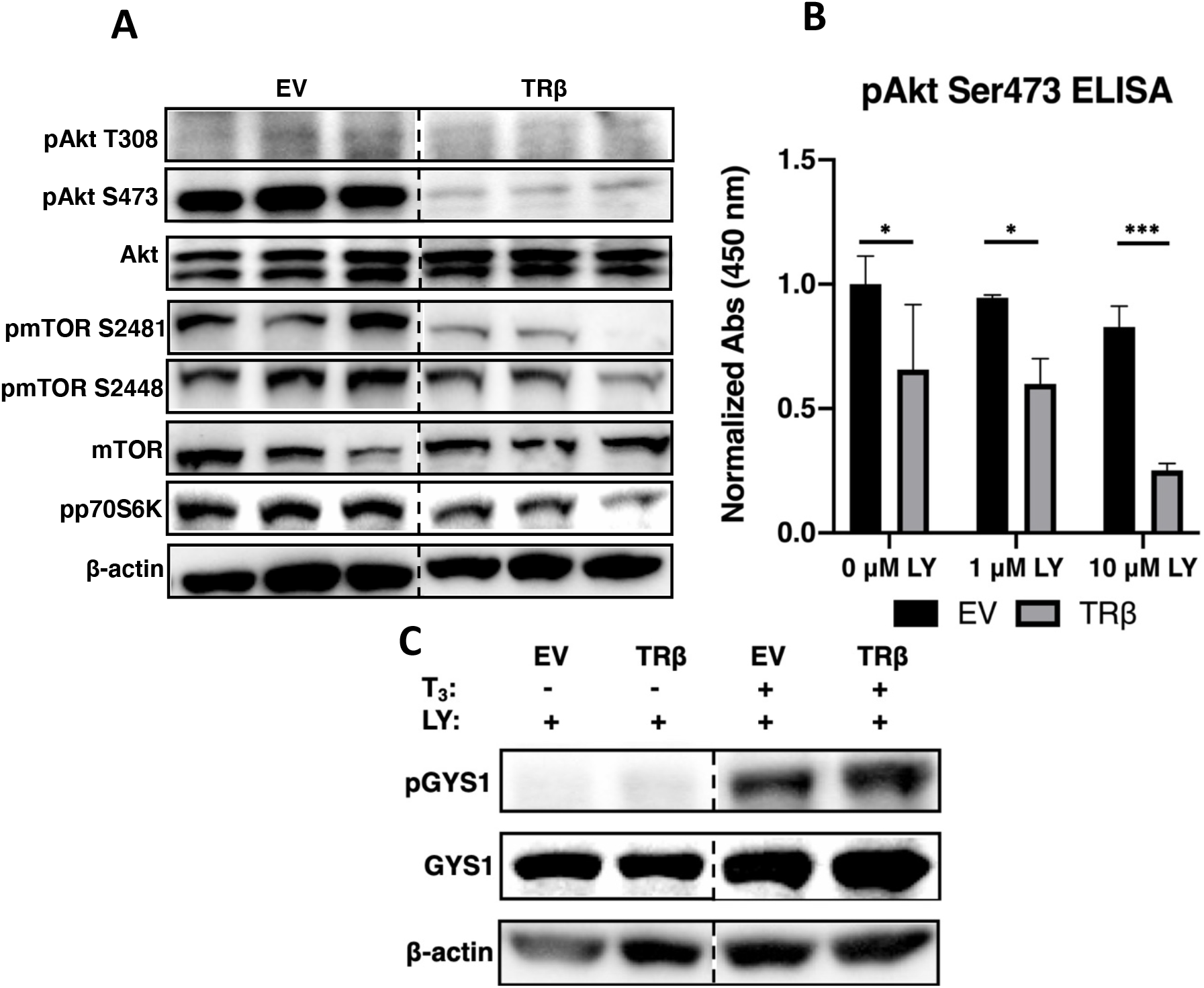
TRβ enhances LY294002-mediated inactivation of the PI3K-Akt-mTOR axis. SW1736-EV (EV) and SW1736-TRβ (TRβ) cells were treated with 10 nM T_3_ for 24 hours before 1 hour of LY294002 (LY) treatment (**A** and **C:** 10 μM, **B:** 0, 1, or 10 μM). Protein levels were determined by immunoblot (**A** and **C**) or subjugated to a pAkt Ser473 sandwich ELISA (**B**). Samples in **A** are replicates. Dashed lines in immunoblots indicate gap between two sets of lanes on the same membrane. ELISA signal in **B** was standardized to protein concentration as determined by a Bradford assay. Significance in **B** was calculated by one-way ANOVA followed by Sidak’s multiple comparison test. (*p < 0.05, ***p < 0.001).

In addition to the Akt-mTOR-p70S6K axis, we also observed an increase in pGYS1 following both T_3_ and LY294002 or buparlisib treatment in our cells, suggesting a robust Akt suppression due to TRβ activation in SW1736 cells (Fig. 8C and Supplemental Fig. 5B). Finally, these results appeared to be dependent on long-term T_3_ treatment, as short-term T_3_ exposure failed to significantly enhance suppression of PI3K with TRβ or LY294002 (Fig. 9A and B).

**Figure 9.**
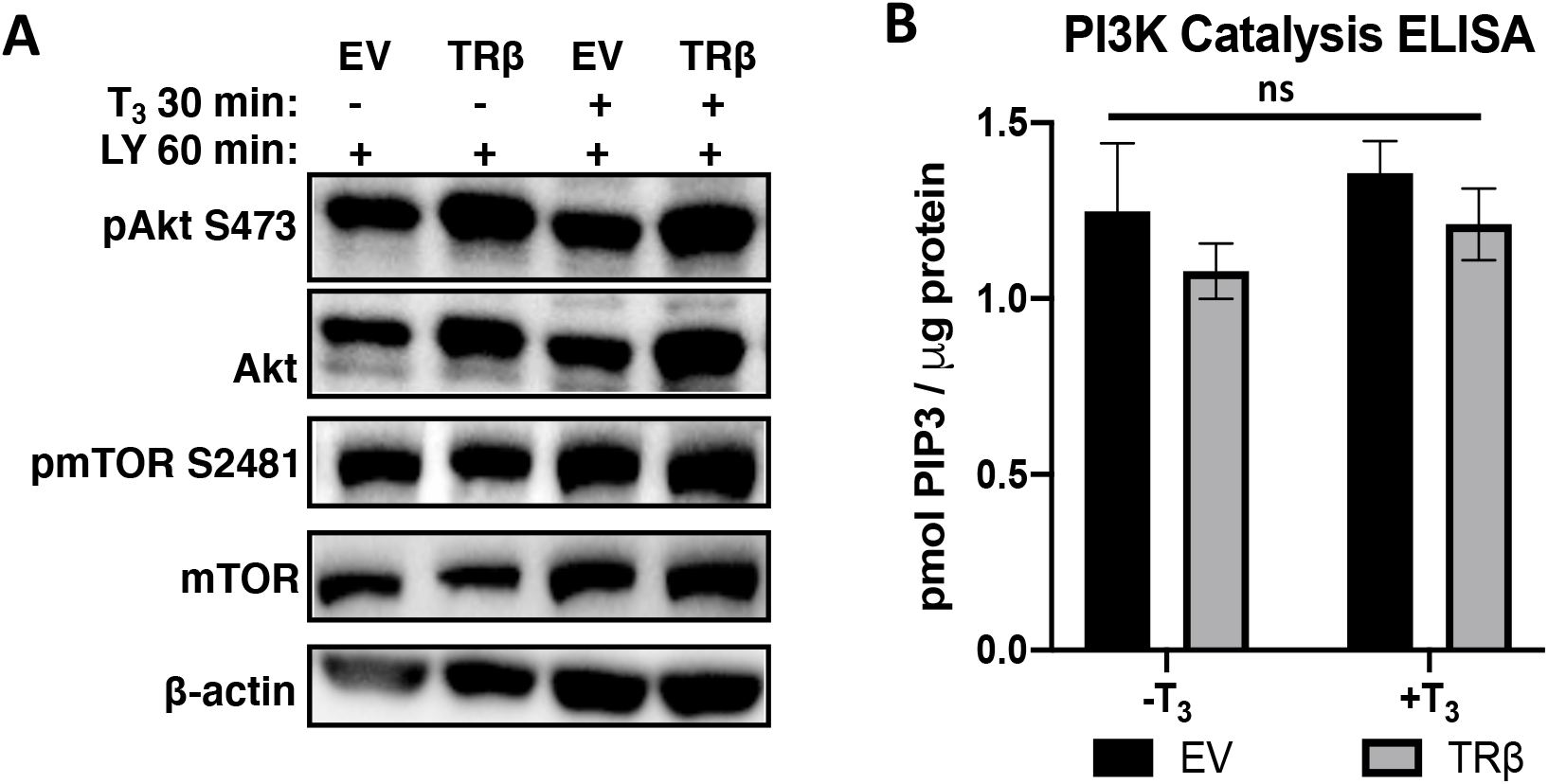
Short-term or no exposure to T_3_ is insufficient for TRβ to enhance LY294002 suppression of PI3K. SW1736-EV (EV) and SW1736-TRβ (TRβ) cells were treated with 10 nM T_3_ or vehicle (1 N NaOH) for 30 min before 1 hour of 10 μM LY294002 (LY) treatment. Protein levels were determined by immunoblot (**A**) or incubated with anti-p85α antibody for PI3K catalysis ELISA (**B**). ELISA signal in **B** was standardized to protein concentration as determined by a Bradford assay. Significance in **B** was calculated by two-way ANOVA followed by Tukey multiple comparisons test. ns = no significance (p ≥ 0.05) across treatment groups.

## 4. Discussion

TRβ has shown to be a potent tumor suppressor in several types of cancer including breast and thyroid cancer (15,16,35,36). However, there are only a few tumor suppressive mechanisms delineated in the literature, which include TRβ-mediated JAK1/STAT1 activation and binding to p85α (15,20,35). Multiple groups have shown the potential for TRβ to bind p85α in the cytoplasm, inhibiting PI3K-mediated PIP_2_ to PIP_3_ catalysis. While TRβ and T_3_ failed to rapidly inhibit PI3K activity in these ATC cells, transcriptomic data revealed novel genomic mechanisms of TRβ-mediated PI3K suppression. Intriguingly, negatively regulated genes included RTKs such as FGFR isoforms and HER3, both of which are implicated in advanced thyroid carcinomas (37–39). TRβ-mediated downregulation of HER3 is particularly interesting, as HER3 is the most potent activating binding partner of HER2 and was found to promote PI3K inhibitor resistance (40,41). In addition to regulating expression of upstream regulators of PI3K, TRβ increased expression of membrane-associated phosphatases, including *INPP5J* and *INPP4B*. *INPP4B* has shown to exhibit remarkable tumor suppression in an *in vivo* model of thyroid cancer by regulating PI3K signaling (42). Furthermore, TRβ also increased expression of the phosphatases *PTPL1*, *PHLPP1*, and the R5B subunit of PPP2. These phosphatases preferentially dephosphorylate ser473 on Akt, and INPP4B and INPP5J modulate ser473 phosphorylation levels (28–30). Increased expression of these phosphatases would account for the robust decrease in ser473 phosphorylation but not thr308 and further supports the notion that TRβ-mediated suppression of the PI3K pathway in these cells is primarily driven by transcriptional regulation. Importantly, both Akt residues must be phosphorylated for maximum activity, as thr308 phosphorylation accounts for only 10% of Akt activity (43). Therefore, the robust pAkt ser473 dephosphorylation may be sufficient to inhibit Akt activity, accounting for the growth and migration inhibition observed in this study. Downstream consequences of TRβ-mediated dephosphorylation of Akt was observed in three well-established downstream targets of Akt, including GYS1, mTOR, and p70S6K.

In addition to the previously documented mechanisms of TRβ tumor suppression, genomic regulation of PI3K regulators likely drives a reduction in cell viability and migration. TRβ enhanced the effect of PI3K inhibitors LY294002 and buparlisib as shown by decreased cell viability, migration, and cell signaling. Importantly, in a phase II trial of ATC patients, buparlisib was able to reduce tumor burden and modestly improve patient survival (44). As we have observed an enhanced response to PI3K inhibitors in the presence of TRβ and hormone, it would be worthwhile to determine if expression levels of TRβ in patient tumors were correlated with patient outcomes.

In summary, our results demonstrate that TRβ suppresses PI3K signaling in SW1736 cells by genomic regulation of RTKs and phosphatases. Although the potential for TRβ to suppress PI3K signaling by binding to the p85α subunit in the cytoplasm is well known, this is the first report to highlight genomic mechanisms by which TRβ suppresses the PI3K-Akt-mTOR axis. The presence and activation of TRβ in ATC cells may be a promising therapeutic target to constrain tumor progression and resistance to chemotherapeutics.

## Abbreviations

Akt: protein kinase B;
ATC: anaplastic thyroid cancer;
AXL: tyrosine-protein kinase receptor UFO;
EGFR: epidermal growth factor receptor;
ELISA: enzyme-linked immunosorbent assay;
FGFR3/4/L1: fibroblast growth factor receptor 3/4/like 1;
FTC: follicular thyroid cancer;
GSK3β: glycogen synthase kinase 3 beta;
GYS1: glycogen synthase 1;
HER2: receptor tyrosine-protein kinase erbB-2;
HER3: receptor tyrosine-protein kinase erbB-3;
INPP4B: inositol polyphosphate 4-phosphatase type II;
INPP5J: phosphatidylinositol 4,5-bisphosphate 5-phosphatase A;
JAK1: janus kinase 1;
mTORC1/2: mammalian target of rapamycin complex 1/2;
p70S6K: ribosomal protein S6 kinase beta-1 PDPK1, 3-phosphoinositide dependent protein kinase 1;
PDTC: poorly differentiated thyroid cancer;
PEKHA2: tandem-PH-domain-containing protein-2;
PHLPP1: PH domain and leucine-rich repeat-protein phosphatase 1;
PI(3,4)P2: phosphatidylinositol 3,4-bisphosphate;
PI(3)P: phosphatidylinositol 3-phosphate;
PI(4,5)P2: phosphatidylinositol 4,5-bisphosphate;
PI3K: phosphoinositide 3 kinase;
PIP3: phosphatidylinositol 3,4,5 trisphosphate;
PPP2R5B: protein phosphatase 2R5B;
PTC: papillary thyroid cancer;
PTEN: phosphatase and tensin homolog;
PTPN13: protein-tyrosine-phosphatase-like protein-1;
ROR1: neurotrophic tyrosine kinase receptor-related 1;
RT-qPCR: reverse transcriptase quantitative polymerase chain reaction;
RTK: receptor tyrosine kinase;
RUNX2: runt-related transcription factor 2;
STAT1: signal transducer and activator of transcription 1;
T3: triiodothyronine;
TRβ: thyroid hormone receptor beta

## Acknowledgments

The SW1736 and KTC-2 cell lines were generously provided by Dr. John Copland III (Mayo Clinic), and 8505C, OCUT-2, and CUTC60 cells were generously provided by Dr. Rebecca Schweppe (University of Colorado). Human cell line authentication, NextGen sequencing, automated DNA sequencing and molecular imaging was performed in the Vermont Integrative Genomics Resource supported by the University of Vermont Cancer Center, Lake Champlain Cancer Research Organization, and the UVM Larner College of Medicine. Additional human cell line authentication was performed by the CU Cancer Center Tissue Culture Shared Resource supported by NCI P30CA046934. Lentiviral constructs were made with the assistance of Dr. Jon Ramsey, University of Vermont (UVM) Cancer Translational Research Laboratory, UVM. Biorender vector graphics software was used to generate Figure 3A. We thank Dr. Jane Lian for early conceptualization and design of this project. We thank Dr. Eyal Amiel for his generous donation of several antibodies and for support on the manuscript discussion.

