## Supplemental Data for "Thyroid hormone receptor beta inhibits the PI3K-Akt-mTOR signaling axis in anaplastic thyroid cancer via genomic mechanisms"

### Supplemental Table 1: Western Blot Antibody Table

| Antigen | Manufacturer; Catalog Number | Species | Dilution |
|---|---|---|---|
| β-actin | Thermo Fisher Scientific; MA5-15739 | Mouse | 1:5000 |
| pAkt (T308) | Cell Signaling Technology; 13038 | Rabbit | 1:1000 |
| pAkt (S473) | Cell Signaling Technology; 4060 | Rabbit | 1:1000 |
| Akt (pan) | Cell Signaling Technology; 2920 | Mouse | 1:1000 |
| pmTOR (S2448) | Cell Signaling Technology; 2971 | Rabbit | 1:1000 |
| pmTOR (S2481) | Cell Signaling Technology; 2974 | Rabbit | 1:1000 |
| mTOR | Cell Signaling Technology; 4517 | Mouse | 1:1000 |
| pp70S6K | Cell Signaling Technology; 9234 | Rabbit | 1:1000 |
| pGYS1 | Millipore Sigma; 07-817 | Rabbit | 1:1000 |
| GYS1 | Abcam; 40810 | Rabbit | 1:1000 |
| TRβ | Millipore Sigma; ABN25 | Rabbit | 1:1000 |
| Mouse IgG | Cell Signaling Technology; 7076 | Horse | 1:10000 |
| Rabbit IgG | Cell Signaling Technology; 7074 | Goat | 1:10000 |

### Supplemental Table 2: Primer Table

| Gene | Forward Oligo (5'->3') | Reverse Oligo (5'->3') |
|---|---|---|
| PTEN | TGTAAAGCTGGAAAGGGACGA | GGAATAGTTACTCCCTTTTTGTCTC |
| INPP5J | CAGGGTGTTGCCCAAACTG | GGCTGCGTTCTTCTTTGCT |
| INPP4B | TGACTGGTGTCACATTCCCAT | GCATAGCCCAAGAAACTTCGC |
| PHLPP1 | GTTCTGCCACTAATTGGTGGA | GCTGGGATGCAACCTTGGA |
| GAPDH | ATGTTCGTCATGGGTGTGAA | TGTGGTCATGAGTCCTTCCA |

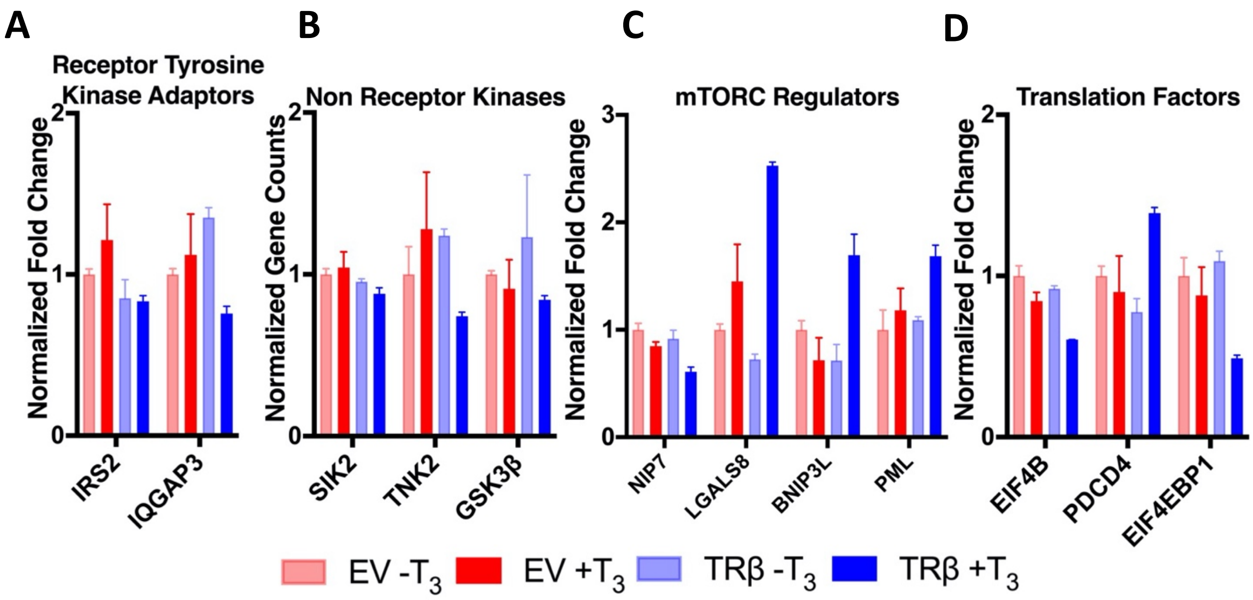


**Supplementary Figure 1: Liganded-TRβ mediates genes involved in PI3K recruitment, PI3K signaling, mTORC stabilization, and translation factors.**

**A.** Ligand-bound TRβ decreased expression of receptor tyrosine kinase adaptors. **B.** Ligand-bound TRβ decreased expression of non receptor kinases involved in amplifying PI3K signaling. **C.** Ligand-bound TRβ regulated expression of mTORC regulators. **D.** Ligand-bound TRβ decreased expression of translation factors downstream of mTORC1. Significance and fold change values between EV+$T_3$ and TRβ+$T_3$ are located in Supplementary Table 1.

| Gene | Protein | Fold Change | Adjusted p-value |
|---|---|---|---|
| ***ERBB3*** | HER3 | 0.787 | 4.03e-6 |
| ***FGFR3*** | FGFR3 | 0.506 | 8.30e-8 |
| ***FGFR4*** | FGFR4 | 0.374 | 1.22e-7 |
| ***FGFRL1*** | FGFRL1 | 0.371 | 8.88e-10 |
| ***ROR1*** | ROR1 | 0.409 | 3.74e-6 |
| ***AXL*** | AXL | 0.393 | 3.26e-5 |
| ***IRS2*** | IRS2 | 0.697 | 2.00e-2 |
| ***PIK3CA*** | P110α | 1.044 | 8.95e-1 |
| ***PIK3CD*** | P110δ | 3.28 | 2.48e-23 |
| ***PIK3R1*** | p85α | 0.770 | 5.92e-2 |
| ***SIK2*** | SIK2 | 0.850 | 3.69e-2 |
| ***IQGAP3*** | IQGAP3 | 0.680 | 2.21e-3 |
| ***PDPK1*** | PDPK1 | 1.036 | 8.04e-1 |
| ***PTEN*** | PTEN | 1.173 | 2.83e-1 |
| ***INPP5J*** | INPP5J | 7.554 | 1.13e-31 |
| ***INPP4B*** | INPP4B | 1.810 | 8.64e-5 |
| ***PLEKHA2*** | TAPP2 | 1.274 | 5.84e-3 |
| ***PTPN13*** | PTPL1 | 1.937 | 2.62e-15 |
| ***TNK2*** | TNK2 | 0.581 | 1.31e-3 |
| ***NIP7*** | NIP7 | 0.719 | 9.28e-4 |
| ***AKT1*** | AKT1 | 1.180 | 2.10e-1 |
| ***AKT2*** | AKT2 | 0.893 | 3.93e-1 |
| ***AKT3*** | AKT3 | 2.307 | 4.10e-1 |
| ***PPP2CA*** | PPP2CA | 1.309 | 8.87e-2 |
| ***PPP2R5B*** | PPP2R5B | 2.530 | 5.19e-9 |
| ***PHLPP1*** | PHLPP1 | 1.513 | 3.90e-5 |
| ***GSK3B*** | GSK3β | 1.586 | 3.41e-6 |
| ***GYS1*** | GYS1 | 1.256 | 1.39e-1 |
| ***BNIP3L*** | NIX | 2.326 | 7.14e-7 |
| ***LGALS8*** | GALECTIN-8 | 1.735 | 9.96e-7 |
| ***PML*** | PML | 1.450 | 1.84e-2 |
| ***MTOR*** | mTOR | 1.005 | 9.77e-1 |
| ***RPS6KB1*** | P70S6K | 1.125 | 6.79e-1 |
| ***EIF4EBP1*** | 4E-BP1 | 0.561 | 5.30e-06 |
| ***EIF4B*** | EIF4B | 0.717 | 7.01e-08 |
| ***PDCD4*** | PDCD4 | 1.532 | 2.99e-3 |

**Supplementary Table 3. Re-expression of TRβ suppresses PI3K signaling genes in SW1736 cells in the presence of $T_3$. Fold change values are relative of TRβ-$T_3$/EV-$T_3$.**

| Cell Line | BRAF Mutation | PI3K Mutation | p53 Mutation |
|---|---|---|---|
| **SW1736** | V600E | N/A | Truncated |
| **8505C** | V600E | N/A | Missense |
| **OCUT2** | V600E | Kinase Domain Gain-of-Function | N/A |
| **CUTC60** | V600E | N/A | Missense |
| **KTC-2** | V600E | N/A | N/A |

**Supplementary Table 4. Cell lines used in Figure 4 (51).**

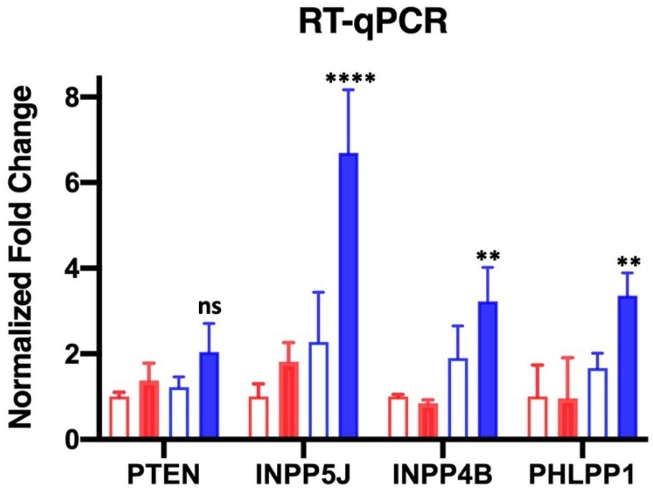


**Supplementary Figure 2: Liganded-TRβ increases expression of inositol and Akt phosphatases.** SW1736-EV (EV) and SW1736-TRβ (TRβ) cells were treated with 10 nM $T_3$ for 24 hours before mRNA levels were determined RT-qPCR. mRNA levels were normalized to the EV-$T_3$ group and significance was calculated by two-way ANOVA followed by Tukey multiple comparisons test (ns = no significance $p \geq 0.05$, $^{**}p < 0.01$, $^{****}p < 0.0001$, significance shown is relative to the EV +$T_3$ group).

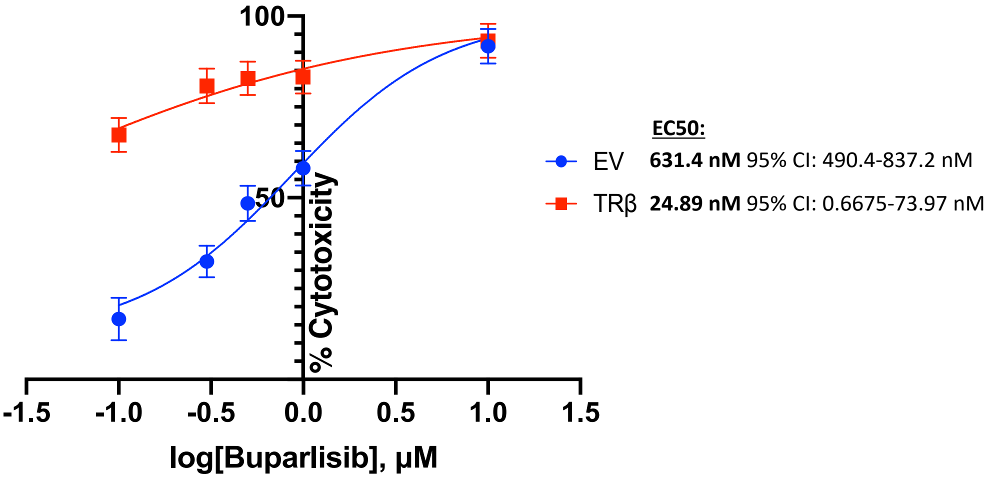


**Supplementary Figure 3: The effect of buparlisib on cytotoxicity is enhanced in SW1736-TRβ cells.**

EV and TRβ cells were treated with 10 nM $T_3$ and buparlisib (0.1-10 µM) or matched-concentration vehicle (DMSO) for four days. Each day the cells were lifted with trypsin and counted using a hemocytometer for viable cells. Area under the curve analysis was conducted for each buparlisib concentration to calculate % cytotoxicity relative to EV vehicle at each concentration of buparlisib. EC50 values were calculated using GraphPad Prism's nonlinear regression package.

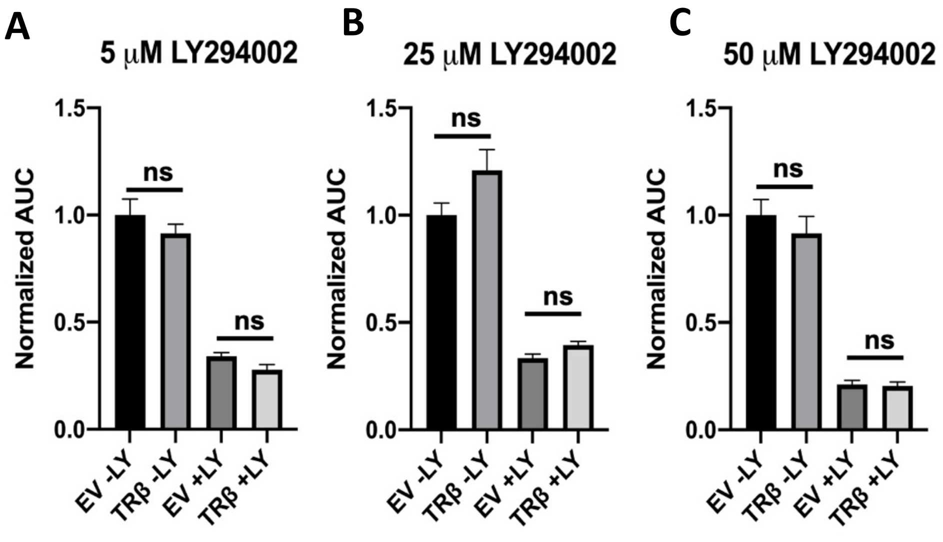


**Supplementary Figure 4: TRβ requires $T_3$ to enhance Ly294002-mediated cytotoxicity.** SW1736-EV (EV) and SW1736-TRβ (TRβ) cells were treated with 1 N NaOH and 5 (**A**)**,** 25 (**B**)**,** or 50 (**C**) µM LY294002 for four days. Each day the cells were lifted with trypsin and counted using a hemocytometer for viable cells. Area under the curve (AUC) analysis was performed and normalized to the EV-LY group. Significance was calculated using one-way ANOVA followed by Tukey's multiple comparison test. ns = no significance ($p \geq 0.05$) across treatment groups.

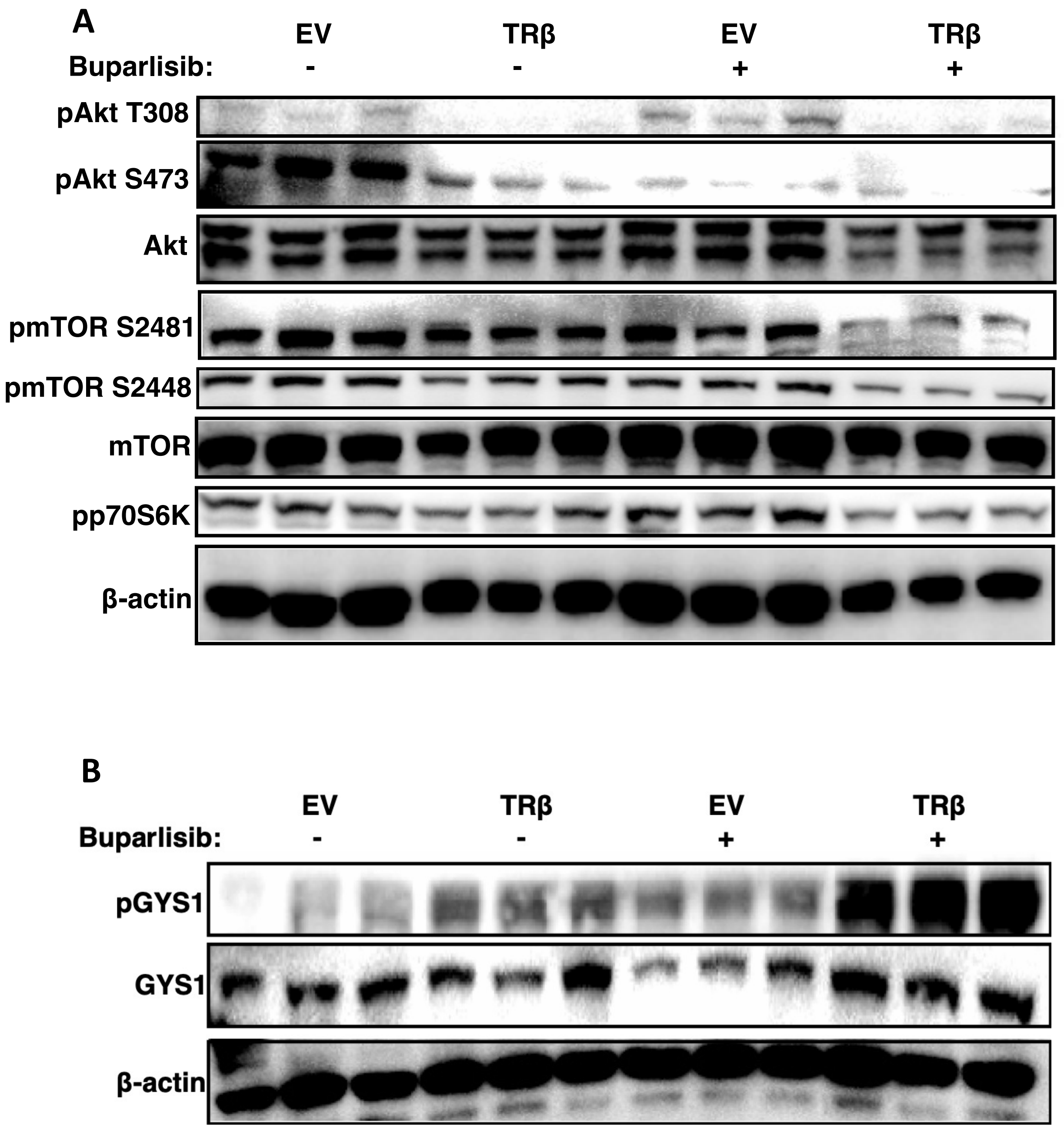


**Supplementary Figure 5: TRβ enhances buparlisib-mediated inactivation of the PI3K-Akt-mTOR axis.**

**A** and **B.** SW1736-EV (EV) and SW1736-TRβ (TRβ) cells were treated with 10 nM $T_3$ for 24 hours before 1 hour of 1 μM buparlisib. Protein levels in triplicate were determined by immunoblot.
